# Adaptive anonymity: Crypsis as an evolutionary trait of floral yeasts?

**DOI:** 10.1101/088179

**Authors:** Moritz Mittelbach, Andrey M. Yurkov, Dominik Begerow

**Affiliations:** Department of Geobotany, Ruhr-University Bochum, Universitaetsstr. 150, 44801 Bochum, Germany; Leibniz Institute DSMZ – German Collection of Microorganisms and Cell Cultures, Inhofferstr. 7B, 38124 Braunschweig, Germany

**Keywords:** Nectar Yeast, Bumblebees, Foraging behavior, *Metschnikowia*, *Cryptococcus*, Pollination

## Abstract

Nectar-dwelling yeast and bacteria are common inhabitants of flowers and evidently involved in pollination. The limited number of floral plant-pollinator models studied to date reveal inconsistent conclusions on microbial effects, but coincide with respect to high microbial specificity: while bacteria reduce visitation frequencies of pollinators, nectar-borne specialist yeasts (in contrast to allochthonous or transient species) impose none or even a beneficial effect on flower visitation. However, these findings are in conflict with the strong impact of these predominantly fermenting organisms on the nectar environment. In order to cope with the ultimate dependency of nectar-dwellers on repeated transportation by foragers as a result of early floral senescence, the modifications of nectar associated with specialist growth have been interpreted as adaptations that suit forager’s preferences. But, the development of foraging preferences to either axenic flowers or flowers colonized by specialist microorganisms would lead to a dead-end for nectar-dwellers, as the probability of inoculation into new suitable habitats would be reduced.

Based on a critical survey of the available literature and an additional pollinator experiment where we find that the allochthonous species *Cryptococcus victoriae* negatively affects attraction and rewarding of floral visitors, while the specialist yeast *Metschnikowia reukaufii* does not, we propose the hypothesis that nectar-borne yeasts may have evolved to blend into their environment avoiding detection by pollinators, following the ecological concept of crypsis. Although, neither chemical nor olfactory crypsis has been reported for nectar-borne microorganisms, the attention to this mechanism in yeast dispersal needs to be directed in future studies.

## 1 The floral niche

The nectar-dwelling community consists of few specialized yeasts (Ascomycota, Saccharomycetales) (Brysch-Herzberg, 2004) and bacteria (Actinobacteria, Firmicutes, and Proteobacteria) (Herrera and Vega, 2012), and a large number of allochthonous species, isolated at low frequencies (Herrera et al., 2010; Junker and Keller, 2015; Mittelbach et al., 2015). While adapted species are primarily vectored between floral habitats by flower visitors, transient species are most probably introduced to nectar from extrinsic sources such as plant surfaces, insects or soil (Fonseca and Inácio, 2006; Good et al., 2014). Their low observation frequencies are believed to result from both random dispersal and from the filtering ability of nectar, which supports a few specialist species such as typical nectar-borne ascomycetous yeasts (Lachance, 2006). However, physiological adaptations to harsh environmental conditions in nectar, such as high osmotic pressure, limited nitrogen supply and plant-derived secondary compounds, might not be sufficient to specialize in this niche (Gruess, 1915). Unlike more persistent plant surfaces that support abundant microbial communities over long time, early floral senescence demands the ability to ensure fast multiplication of cells and the repeated dispersal of propagules into new floral habitats. This degree of dependency on inter-floral dispersal by pollination vectors (e.g. insects or birds) should be higher for specialist nectar-dwellers (autochthonous) than for widespread (sometimes also erroneously referred to as ubiquitous) allochthonous species. The latter are not restricted to floral habitats and are called generalists (as opposed to true nectar yeasts) in the ecological literature (Vannette et al., 2013). Fundamental assumptions regarding the dispersal of specialist and generalist species are supported by the fact that well-known nectar-dwelling species are seldom isolated from sources other than nectar or related pollinators (Brysch-Herzberg, 2004; Fonseca and Inácio, 2006; Lachance, 2006). Independent of the degree of specialization in the floral niche, proliferation of microorganisms leads to chemical alterations of the nectar that, in turn, might interfere in tightly linked interactions of plants and pollinators. Most likely, nectar-borne microorganisms may influence two crucial and costly mechanisms of pollination, namely the attraction of pollinators to flowers through olfactory cues, and the subsequent rewarding of visitors via the provision of nectar. Although evidence is still lacking in the scientific literature, plant-emitted olfactory signals could be altered by microbial intervention (Golonka et al., 2014) in a way that affect pollinators’ decisions to visit a certain flower (Pozo et al., 2009, Schaeffer et al., 2016). Also, microbially derived alterations of the nectar itself, such as reduced nutritional value (sugar depletion), increased acidity (Vannette et al., 2013) and ethanol contents (Mittelbach et al., 2016), may not fit well foragers preferences and lead to reward-based decisions affecting the subsequent visitation (Nepi, 2014).

One can assume that the dependency of specialized nectar-borne yeasts on repeated transportation by pollinators presumes scarcely any perceptible impact on the habitat itself in order to maintain dispersal pathways. Thus, from an evolutionary point of view, growth of nectar-associated yeasts is expected to be either negligible (neutral effect) or to meet the expectations of pollinators (positive effect). In contrast to true nectar-borne yeasts, widespread generalists often possess a broader enzymatic machinery, which allows them to consume a wide range of carbon sources and therefore, to survive on both living and dead plant material (Fonseca and Inácio, 2006). They do not necessarily rely on further dispersal by pollinators and can compromise impoverished pollination services. Thus, obviously different lifestyles of nectar-dwelling specialists and generalists should be considered in modeling interactions of microorganisms with plant-pollinator systems.

## 2 Microbial imprint on pollinator foraging

Considering current knowledge of yeast-pollinator interactions, direct effects of microbial nectar-dwellers on mechanisms of pollination and plant fecundity are so far contradictory (Table 1). Experiments in flight cages with artificial flowers (Herrera et al., 2013, Schaeffer et al., 2016), and in portable glasshouses with natural flowers (Schaeffer et al., 2014) indicate that bumblebees prefer nectar colonized by one of the most frequent nectar-borne specialist yeasts *Metschnikowia reukaufii* over artificial media or sterile nectar, respectively. Yeast inoculation enhanced visitation frequencies, but did not to change the foraging time in a single flower, so the observed effects have been explained by the alteration of floral attraction, without a negative impact on rewarding (Schaeffer et al., 2014). In another study both attraction of bumblebees and nectar consumption in flowers of *Hellerborus foetidus* were positively affected by the growth of *M. reukaufii* (Herrera et al., 2013). A few studies demonstrated that unlike bumblebees, honeybees and hummingbirds avoid nectars containing prokaryotic microorganisms, but showed no response to *M. reukaufii* (Good et al., 2014; Kevan et al., 1988; Vannette et al., 2013). In summary, the aforementioned studies show a high degree of context-dependency in pollinator foraging behaviour: pollinators confidently preferred yeast-inoculated nectars only under controlled conditions (e.g. in flight cages) and without an alternative treatment offering a nectar-borne competitor. When other microorganisms were alternatively offered in experimental trials or if distraction (here: the diversity of the nectar-borne meta-community within the foraging range of the pollinator) was presumably high, pollinators did not respond to the presence of *M. reukaufii* in flowers.

**Table 1.**
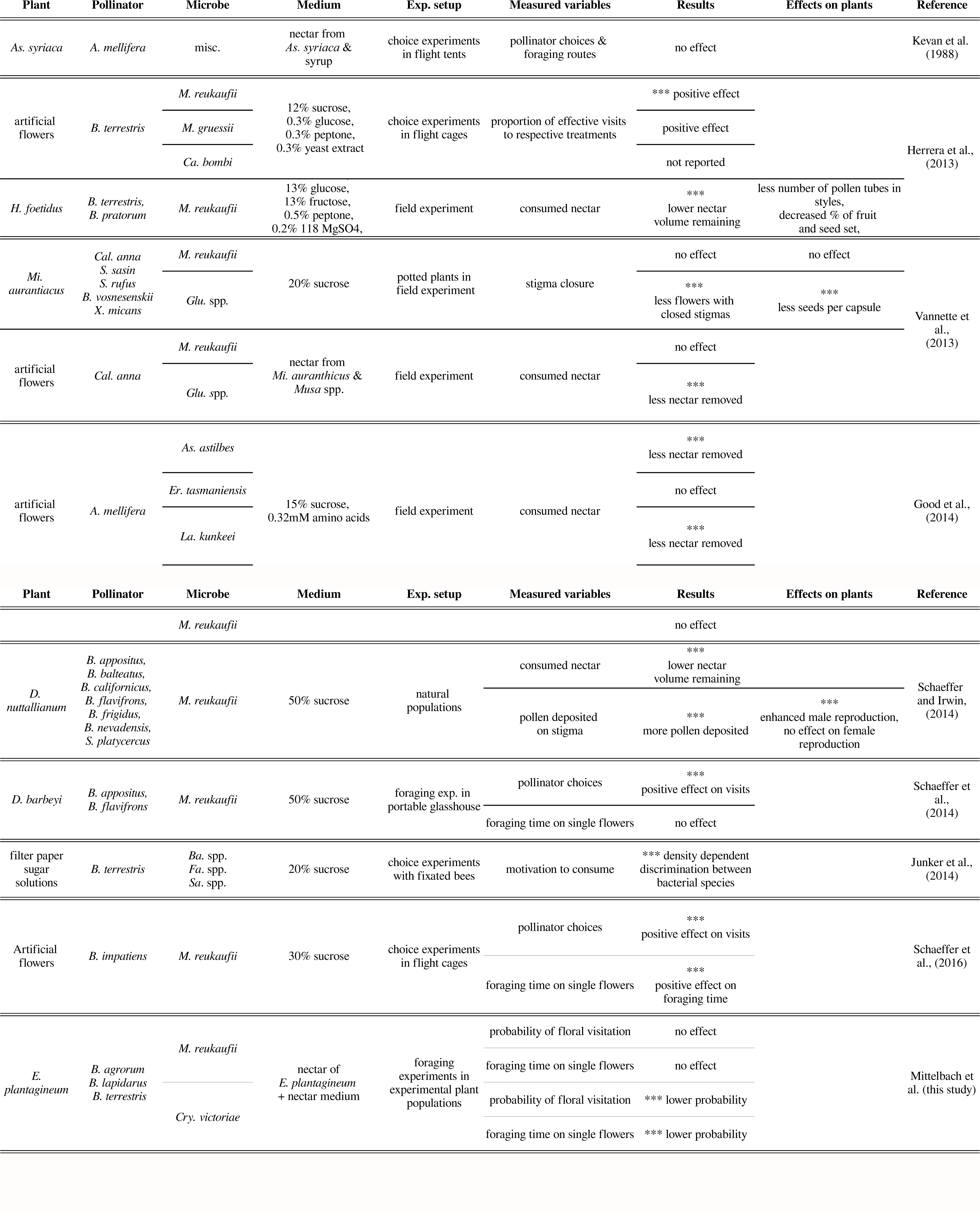
Summary of published results on effects of nectar-borne microbial organisms on pollination and pollinator foraging. Only significant results (*p <= 0.05*) are considered as effect. Species names are abbreviated: Bacteria: (As) *Asaia* and (Er) *Erwinia* (both Gram negative bacteria), (Ba) *Bacillus* (Fa) *Falsibacillus* (La) *Lactobaccillus* (Sa) *Salirhabdus* (Gram positive bacterium), (Glu) *Gluconobacter* (Proteobacteria); Fungi: (M) Metschnikowia, (Ca) Candida (both Ascomycota), (Cry) *Cryptococcus* (Basidiomycota); Insects: (B) *Bombus*, (A) *Apis*, (X) *Xylocopa* (all Apidae); Birds: (S) *Selasphorous*, (Cal) *Calypte* (both Trochilidae); Plants: (H) *Helleborus* and (D) *Delphinium* (both Ranunculaceae), (Mi) *Mimulus* (Scrophulariaceae), (As) *Asclepias* (Apocynaceae), (E) *Echium* (Boraginaceae).

## 3 Crypsis as adaptation of nectar-borne specialists

Taken together, studies that collectively analyzed nectar removal, pollinator choices, pollen transfer, and stigma closure (Table 1) did not provide a solid hypothesis on microbial adaptation to pollination, but agreed on the absence of negative effects by nectar specialist yeasts on foraging strategies of floral visitors. The observed behavioral changes of foragers seem to be a product of focal nectar-borne species and actual alternative choices presented to pollinators. Therefore, we propose the hypotheses that nectar-borne specialists might have taken a different way to ensure the dispersal by pollinators through developing chemical and olfactory crypsis. In ecology, crypsis is the ability of an organism to avoid detection or identification by another organism in environmental contexts (Ruxton, 2009). This adaptive strategy includes camouflage, mimicry and a range of non-visual (e.g. sound, olfactory, chemical) adaptations that help organisms to avoid predation in respect to defined environmental conditions. Consequently, crypsis depends on defined environmental conditions and might be abolished in changing environments. Although, not yet been applied to microorganisms, in our opinion, the concept chemical crypsis explains well previously reported observations from yeast inoculation experiments, when experimental and environmental conditions are taken into account. The fact that bumblebees are per se able to detect *M. reukaufii* in flowers (Schaeffer et al., 2016) is no contradiction to crypsis theory, but rather a support for its context-dependency. Since, it is unlikely (except perhaps for early-blooming flowers with high sugar contents) that pollinators only have to choose between sterile flowers and flowers colonized by *M. reukaufii*, a nectar-borne species remains unrecognized if forager behaviour is primarily steered by external (negative or positive) stimuli.

Hypothetically, a foraging pollinator in its everyday life needs to discriminate constantly flowers from different plant species, some of which may also harbor different microbial communities. In the same manner as pollinators develop a foraging preference to certain plant species or scents (McAulay et al., 2015), they may choose certain microbial communities above others. Thus, the current local choice, either of plants or microbial communities, always steers foraging decisions. The development of foraging preferences by pollinators to microbial communities would result in negative effects for nectar-borne microbial species, which rely on the constant cycle of inoculation, growth, removal, and transportation during the flowering season. On the one hand, the repellence of pollinators by nectar-dwellers would immediately reduce the dispersal-rate of these microbes. On the other hand, the development of foraging preferences towards colonized flowers would reduce the habitats of species in size because visitation rates to axenic flowers would be reduced and already colonized flowers bear strong competition for late arriving microbial species (Peay et al., 2012). Consequently, both preferences would put nectar yeasts at the risk of extinction from habitat fragmentation or even loss. This mechanisms are not equally true for allochthonous or transient species, which are able to survive and proliferate as saprophytes on other organic or inorganic substrates and the negative impact on pollinator foraging of bacteria and transient yeast species (Table 1) should not have a substantial impact on population development of these species. The unpredictable modifications of the nectar itself by these allochthonous species (Mittelbach et al., 2016) could impede the development of late arriving specialists (Peay et al., 2012), therefore, nectar specialists most likely even benefit from pollinator’s distinctions between autochthonous and allochthonous communities. Based on this assumption, we propose that nectar specialists adapted to the floral niche by continuously reducing their impact on pollinator foraging in order to blend in the environment. This theory also receives support in the recent studies (Table 1), which showed that pollinators discriminated between sterile flowers and flowers colonized by autochthonous species only when no second non-specialist (allochthonous) microbial species was offered.

The proposed concept of crypsis is no exclusive mode in the interaction between nectar-borne yeasts and bumblebees, because detection of yeasts should always depend on the given environmental context and the competing microbial communities present in the local environment. Since bees can principally detect *M. reukaufii* in flowers, preferential visits of sterile flowers might be an experimental artifact when pollinators target non-manipulated flowers for a better nutrition as it is not yet depleted by nectar dwellers (e.g. Herrera et al., 2008). Similar mechanisms may underlie the results obtained for floral systems in extreme environments, such as winter-blooming plants (Herrera et al., 2013). Specifically, both high sugar concentration and early seasonal flowering of plants result in a lower diversity and spatial heterogeneity of nectar-borne microbial communities, as compared to other studied floral systems (Brysch-Herzberg, 2004; Herrera et al., 2008). So the increased removal of nectar in flowers of *H. foetidus* inoculated with *M. reukaufii* indirectly illustrates our theory of external distraction, as the lack of additional choices is likely to rule out foragers’ distraction from flowers and leads to the observed preferential visitation of flowers inoculated with *M. reukaufii* by bumblebees.

As already mentioned, specialist yeasts grow well in co-cultures with other nectar-borne species and often successfully over-compete them in laboratory experiments. Thus, typical nectar specialists may not need to provide a positive effect on attraction and reward of pollinators in order avoid a foraging preference. When a pollinator randomly transfers yeast inoculum to new flowers, the specialist yeast can most likely out-compete already growing generalist species through faster growth, osmotolerance, and competitiveness.

Both pollinator foraging behavior and physiological properties of typical nectar yeasts successfully blending in the environment (crypsis) ensure non-selective random cell transfer between inoculated flowers and flowers not visited before. This hypothetical scenario would result in an equal visitation frequency of flowers inoculated with specialist species and axenic flowers by a pollinator. To gather additional experimental evidence for our hypotheses we performed a field experiment where we manipulated flowers of *Echium plantagineum* with yeast cultures, either of the ascomycetous specialist *M. reukaufii*, the basidiomycetous transient *Cryptococcus victoriae*, or pure growth medium as control (Supplementary). These treatments were offered to foraging bumblebees simultaneously in different flowers of the same inflorescence, and we recorded decisions of individuals to visit a certain flower and the time they spent in visited flowers. We found experimental evidence supporting our hypothesis that *M. reukaufii* does not influence pollinator decisions or the time spent on foraging nectar (Figure 1) compared to pure growth medium, when offered together with a transient species. In agreement to the generalist-specialist hypothesis, the inoculation of *C. victoriae* into flowers weakens the relation between flowers and pollinators and reduces the probability to visit a certain flower and the time spent inside single flowers.

**Figure 1.**
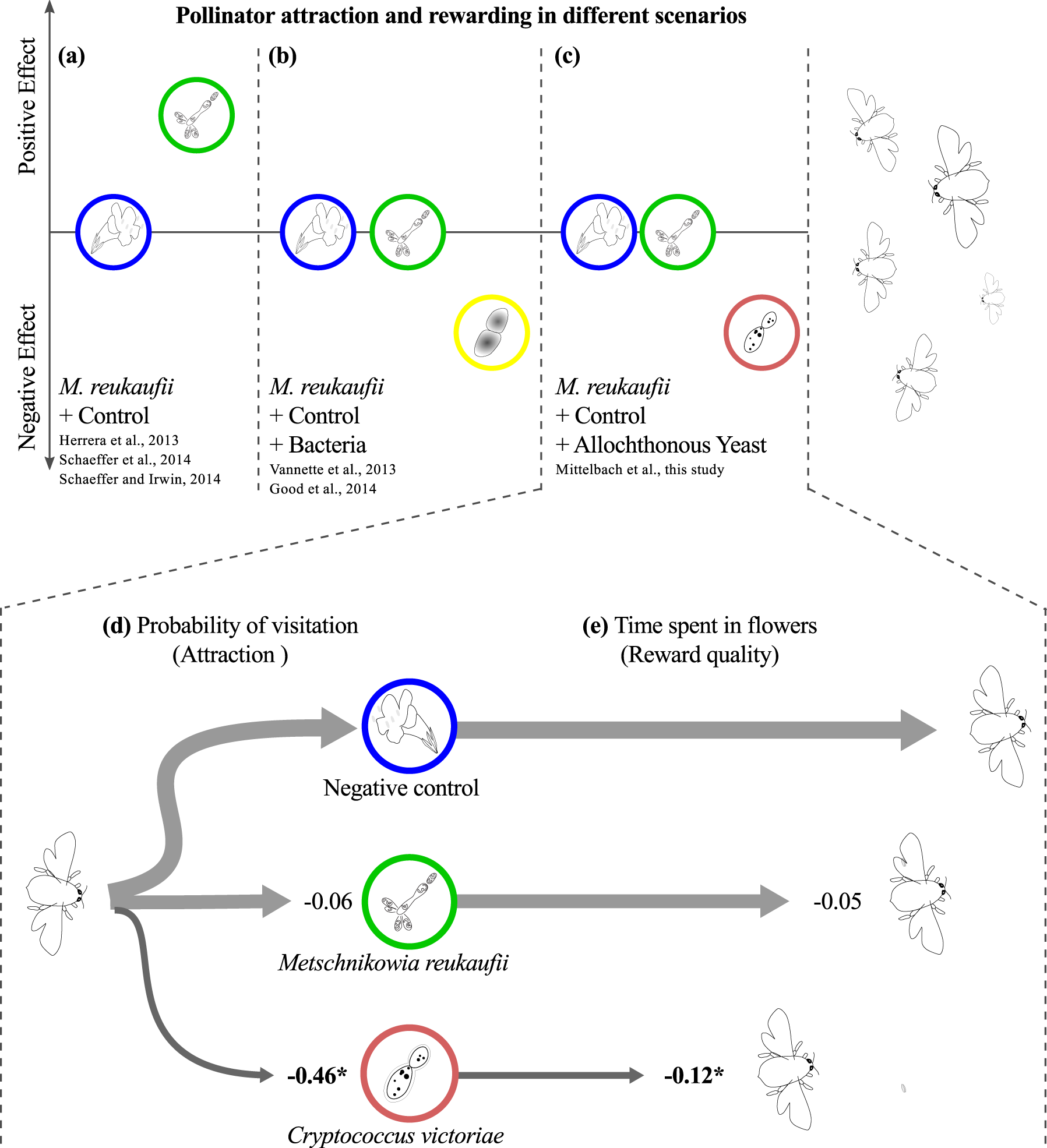
(A) Schematic diagram of foraging responses of pollinators to microbial growth in floral nectar, based on available literature (a,b) and own foraging experiments (c-f). The baseline is set to the response to control flowers (either inoculated with sterile medium or not manipulated) in each of three scenarios, respectively. The first section (a) shows the choice between controls (blue) and flowers inoculated with *Metschnikowia reukaufii* (green). In the second (b) and third (c) scenario either bacteria (yellow) or allochthonous yeasts (red) are added as choice to foragers. (B) Estimated effects in pollinator experiments of inoculation of yeast species (specialist species: *M. reukaufii*, transient species: *Cryptococcus victoriae*) on foraging behavior of bumblebees in flowers of *Echium plantagineum*. We show the probability of floral visitation (d) and the time spent in flowers (e) calculated using hurdle GLMMs. Wald’s z statistics are shown in bold (*: p<= 0.05).

In our opinion pollination is not well explained by yeast attraction of the foragers. The literature survey and our own results suggest that the influence of nectar-borne yeast on pollination is context-dependent and varies in relation to pollination syndrome (Mittelbach et al., 2015), environmental conditions, available choices, and the actual meta-community of microorganisms. Among beneficial mechanisms in pollination scenarios with uncommon conditions, such as temperature increase in winter-blooming plants (Herrera and Pozo, 2010), or notable accumulation of alcohols (Wiens et al., 2008), the main evolutionary advantage of nectar-dwelling ascomycetous yeasts might be their minimal disturbance of the existing interactions between plant-pollinators and the flower, as compared to transient or generalist species. Indeed, our observations provide the first experimental evidence that nectar “contamination” by widespread phylloplane-related yeasts may have a negative effect on plant-pollinator interactions. In our experiment, this negative effect on bumblebees was sufficiently strong to allow *M. reukaufii* to blend into the environment, avoiding the differentiation to sterile flowers by foraging individuals.

This result is in agreement with the hypothesis of reduced niche-reliance of transient species in contrast to specialist nectar-borne species (Vannette et al., 2013). However, we additionally point out that evaluation of microbial impacts on pollination require more comprehensive experimental setups, like the one used in this study. Field experiments should take into account respective characteristics of pollination syndromes, environmental conditions and microbial metacommunities in flowers. Since the cryptic behaviour of nectar yeasts is context-dependent, future studies should evaluate the consequences for yeasts (both specialists and generalists) and pollinators in different scenarios, ranging from random pollinator foraging (complete crypsis) to highly preferential foraging on yeast colonized flowers (absence of crypsis).

In our opinion, the ecological concept of crypsis might be an overlooked mechanism in further microbial mutualistic and parasitic systems, where interactions between trophic levels are responsible for species survival or ecosystem functioning.

## 4 Materials and Methods

### 4.1 Literature survey

We included all available studies, testing the impact of microbial nectar-dwelling species on pollinator foraging, by thoroughly search scientific databases and using search engines (ISI Web of Knowledge, Google Scholar).

### 4.2 Experimental procedure

The nectar-borne ascomycetous (Saccharomycetes, Saccharomycotina) yeast *Metschnikowia reukaufii* Pitt & M.W. Mill. (Strain MOM_296 = DSM 29087) was selected as a representative specialist yeast species (Table 1). Widespread basidiomycetous (Tremellomycetes, Agaricomycotina) yeast *Cryptococcus victoriae* M.J. Montes, Belloch, Galiana, M.D. García, C. Andrés, S. Ferrer, Torr.-Rodr. & J. Guinea (Strain MOM_325 = DSM 29088) was selected as a representative of generalist species based on available reports in the literature (Fonseca et al., 2011; Yurkov et al., 2015). Both strains have been isolated from floral nectar of *Echium plantagineum* L. (Boraginaceae) in the botanical garden of the University Bonn (Germany) in summer 2012.

We recorded 20 independent foraging trials of individual female worker bumblebees, each on a separate plant individual in an experimental population of *E. plantagineum* at the botanical garden of the Ruhr-University Bochum, Germany. To ensure an undisturbed foraging behavior of insect metacommunity, the population consisted of 40 plant individuals, from which 20 individuals were chosen for the use throughout the experiment and covered simultaneously during the experimental trials. The experiment was replicated 4 times (n=80) on 4 days. The main inflorescences of the 20 randomly selected plant individuals were prepared in advance as follows: All cymes insisting at the focal branch were marked with different colored rings. Beginning from the uppermost cyme, colors were assigned to all cymes alternately, following the convolute architecture of the inflorescences. The most upper color mark was chosen randomly. To avoid a systematic effect of the cyme position on bumblebee foraging strategies in inflorescence architectures, we randomly exchanged assignments of inoculations to color codes in each replicate, so that all experimental treatments were presented to foragers in different heights and positions in inflorescences.

At the beginning of each trial of the experiment, all open flowers of focal branches were removed carefully. To prevent bee foraging on buds, we covered the inflorescences prepared for the experiment with sterile gauzes. After 24 hours, we carefully added 0.5 μL of sterile nectar media (1 × YNB + 30% sugar w/l: 70% sucrose, 16% glucose, 14% Fructose) to nectar droplets of all newly opened flowers and flowers in enhanced bud stages (average number of flowers per inflorescence: 15, range: 11-26). Inoculations were assigned consequently to color codes within individual plants and days and contained approx. 1000 cells (2000 cells/μL) of either *M. reukaufi* i or *C. victoriae*, or were yeast-free as negative control. Droplets of the inoculation substances were carefully placed with a laboratory pipette at the bottom of corolla tubes below the base of the central anther where naturally secreted nectar droplets are naturally presented. After the inoculations, branches were carefully covered again for 24 hours. After assuring that bumblebees were actively foraging at uncovered plants in the experimental population, the first gauze was removed. The foraging trial of the first visitor was recorded using a regular video camera (Nikon P100,). Visits by honeybees or interrupted foraging trials were discarded and plants were excluded from data analysis. One foraging trial was defined as all floral interactions of one individual bumblebee at the focal inflorescence, beginning with the first visit or rejection. The trial ended when the observed individual probed at a flower of another inflorescence or left the population entirely. In order to quantify the bout-length and probabilities of visits, the videos were inspected visually. To assure successful colonization by inoculated yeasts and to evaluate probable contaminations, we randomly choose 15 flowers (5 of each treatment) after each experimental replicate and plated the nectar on agar plates containing yeast media (0.3% w/v Yeast extract, 0.5% w/v Peptone, 0.3% w/v Malt extract, 1% w/v Glucose, 1% w/v Fructose and 1% w/v Sucrose, 2% w/v Agar) following a standard protocol (Mittelbach et al., 2015). All plates showed positive growth of the expected species and visual estimations of colony forming units (CFU) revealed only minor contaminations (5 of 60 control plates of flowers inoculated with or without yeast, and max 3 colonies in sampled flowers). Statistical procedures are explained in the Supplementaries.

## Acknowledgments

The authors thank M. Pennekamp for assistance during the experiments, T. Eltz for critical discussion, and M.-A. Lachance and anoymous reviewers for valuable comments on the manuscript. Special thanks go to T. Stützel and the botanical garden of the Ruhr-University Bochum. M. Nepi and D. Nocentini kindly provided data on nectar chemistry of *E. plantagineum*.

## 5 Author contributions

MM planned, conducted and analyzed the experiment with consultancy of DB. The hypothesis was developed by MM and AY. MM wrote the manuscript with assistance of AY and DB.

